# Genetic modification of intractable staphylococcal clones by heat-shock facilitated phage transduction

**DOI:** 10.1101/2025.04.04.647181

**Authors:** Lukas Schulze, Jens Moessner, Sophia Krauss, Theresa Harbig, Kay Nieselt, Bernhard Krismer, Andreas Peschel

**Affiliations:** Department of Infection Biology, Interfaculty Institute of Microbiology and Infection Medicine, University of Tübingen, Tübingen, Germany; Cluster of Excellence EXC 2124 Controlling Microbes to Fight Infections; German Center for Infection Research (DZIF), partner site Tübingen, Germany; Institute for Bioinformatics and Medical Informatics, University of Tübingen, Germany

**Keywords:** Bacteriophages, Staphylococci, Transduction, Restriction-Modification, Restriction barrier, Genetic manipulation, Heatshock

## Abstract

Increasing recognition of commensal bacteria as a requirement for microbiome integrity and pathogen exclusion puts urgency to the molecular characterization of commensal interactions. However, many commensals cannot be transformed with available methodology because of potent restriction barriers. We developed a new method for the introduction of plasmid DNA into otherwise intractable non-*Staphylococcus aureus* (NAS) staphylococci, important commensal members of human nose and skin microbiomes, by phage transduction. We demonstrate that a precise pulse of recipient bacteria with elevated temperatures prior to exposure to transducing phages renders NAS isolates effectively and transiently amenable to transduction. Transduction of NAS mutants lacking restriction-modification (RM) systems did not profit from a heat shock indicating that it is the transient deactivation of RM enzymes that permit transduction. Our method also facilitated the transduction of other Bacillota from the genera *Bacillus* and *Listeria* indicating that it will support research on different bacterial groups from diverse ecosystems.

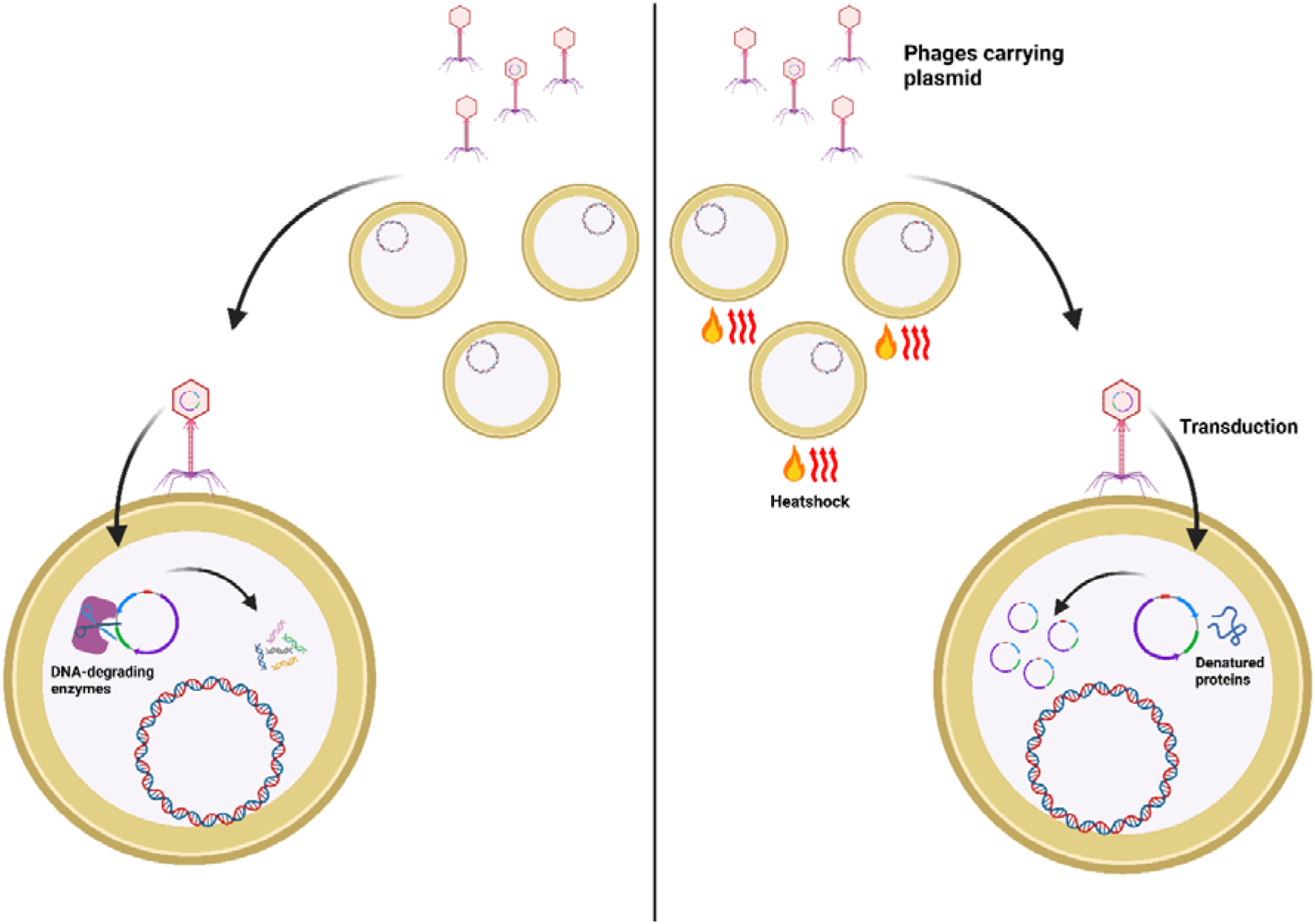

## Introduction

Staphylococci are prominent members of the human microbiome, including both, commensals and facultative pathogens, that impact on health or disease of their host in multiple ways ^1–3^. Much attention has been devoted to the coagulase-positive species *Staphylococcus aureus*, which is an opportunistic pathogen, responsible for severe soft tissue infections, bacteraemia, endocarditis, and many other types of infection ^4,5^. In contrast, infections caused by non-*S. aureus* (NAS) staphylococci are usually less severe, but they have gained increased attention in recent years, because NAS such as *Staphylococcus epidermidis* are major causes of catheter or prosthetic joint-associated infections ^6,7^. Ob the other hand, many isolates from NAS species such as *S. epidermidis, Staphylococcus lugdunensis,* or *Staphylococcus capitis* have also been reported to protect their host from *S. aureus* colonization by the production of *S. aureus* -eliminating antimicrobial secondary metabolites such as bacteriocins ^8–11^. Some bacteriocins have even been found to amplify the immune response or to act synergistically with host-derived antimicrobial peptides, demonstrating the fine-tuned interplay between the human host and its staphylococcal commensals ^12,13^.

Genetic tractability is a prerequisite for studying the lifestyle and the phenotypic traits of bacteria, including those of staphylococci. To this end, a plethora of methods for bacterial genetic transformation and manipulation has been established throughout the last decades. While chemical transformation and conjugation are favourable methods for transferring genetic material into many bacterial species ^14–17^, such methods are not applicable to staphylococci, as natural competence is usually not observed and conjugation is rare in this genus ^18,19^. However, methods for phage-mediated DNA transfer, known as transduction ^18,19^, as well as the development of protocols for transformation by electroporation ^20^ have greatly advanced the ability to study and work with many staphylococcal strains.

Nevertheless, studying staphylococci remains challenging, as especially NAS possess strong and diverse barriers for the introduction of foreign DNA, which impede genetic manipulation and render many strains entirely inaccessible to molecular research. Bacteria can encode four different types of restriction/modification (RM) systems, Type-I, II, III or IV. These systems consist of various combinations of restriction endonucleases (REase), DNA-methyltransferases (MTase), and sequence specificity-conferring proteins. Type-I systems consist of all three proteins while the Type-II and Type-III systems consist only of REase and MTase, and Type-IV systems of just a single REase^21–24^. Previously, all four different RM types have been described in staphylococci ^25–29^. Importantly, individual strains of the same species often express multiple different RM-systems, with different sequence specificities^30^. In addition to RM systems, staphylococci, can express Clustered regularly interspaced short palindromic repeat (CRISPR)-Cas systems, a sophisticated form of bacterial adaptive immunity, for the detection and degradation of foreign DNA ^31,32^.

To circumvent these DNA immunity systems and prevent the degradation of foreign DNA by staphylococcal recipient strains, various strategies have already been published. For example, a short heat shock prior to electroporation significantly increased the transformation efficiency in *S. aureus* and *Staphylococcus carnosus*, probably by denaturing RM proteins transiently ^33, 34^. As this methodology has only been validated for those two strains, a more general procedure, based on plasmid artificial modification (PAM) has been developed. Here, transformation efficiency of recipient strains with a strong RM barrier is enhanced by passaging the plasmid through a specifically engineered *E. coli* host. This intermediary host expresses the RM system-specific methylase to modify the transferred DNA with a suitable methylation pattern that protects it from degradation by the recipient strain ^35,36^. However, this approach is time and labour-intensive, as the modification system of the strain of interest must be identified, cloned, and expressed in *E. coli*. Subsequently, the methylated plasmid must be isolated from this modified strain for electroporation into the staphylococcal strain of interest. Moreover, this approach is usually not efficient for recipients expressing more than one RM system and therefore not suitable for most NAS isolates, which usually have more than one RM system.

We describe a new technique, which combines heat shock and transduction by a suitable phage, to enable the genetic manipulation of previously intractable staphylococcal strains and species. Our experimental data suggests that temporary inactivation of the restriction systems by the heat shock enables the successful introduction of foreign DNA into these strains. The new method is applicable for different plasmids, bacterial species, and transducing phages and it is also effective in non-staphylococcal genera such as *Bacillus* or *Listeria,* demonstrating its potential benefit for the wider microbiological community.

## Results

### Identification of systems protecting from invading DNA

Genetic transformation of colonizing or infecting *Staphylococcus* isolates remains a major obstacle, mostly because of barriers that prohibit the introduction of recombinant DNA via transformation or transduction. We selected the two *S. epidermidis*strains 17-20 and D2-30, isolated from human nasal microbiomes, and the *S. pseudintermedius* strain ED99 from canine skin infection ^37^, which could not be transformed or transduced with standard methods, to elucidate more effective ways for the introduction of plasmid DNA. The genomes of the three strains were analysed for the presence of potential RM and CRISPR Cas systems using the ‘Prokaryotic Antiviral Defence LOCator’ (PADLOC) ^38^. *S. epidermidis* 17-20 encoded two RM systems (Figure 1 A), a Type-I and a Type-II RM system, composed of three and two genes, respectively. The genes of both systems were analysed with the REBASE database ^39^, to identify potential DNA recognition motives. The Type-I and Type-II systems were predicted to use the recognition sequences ‘GAGN_7_TAC’ or ‘GWAGN_6_TTTA’ and ‘GATC’, respectively.

**Figure 1:**
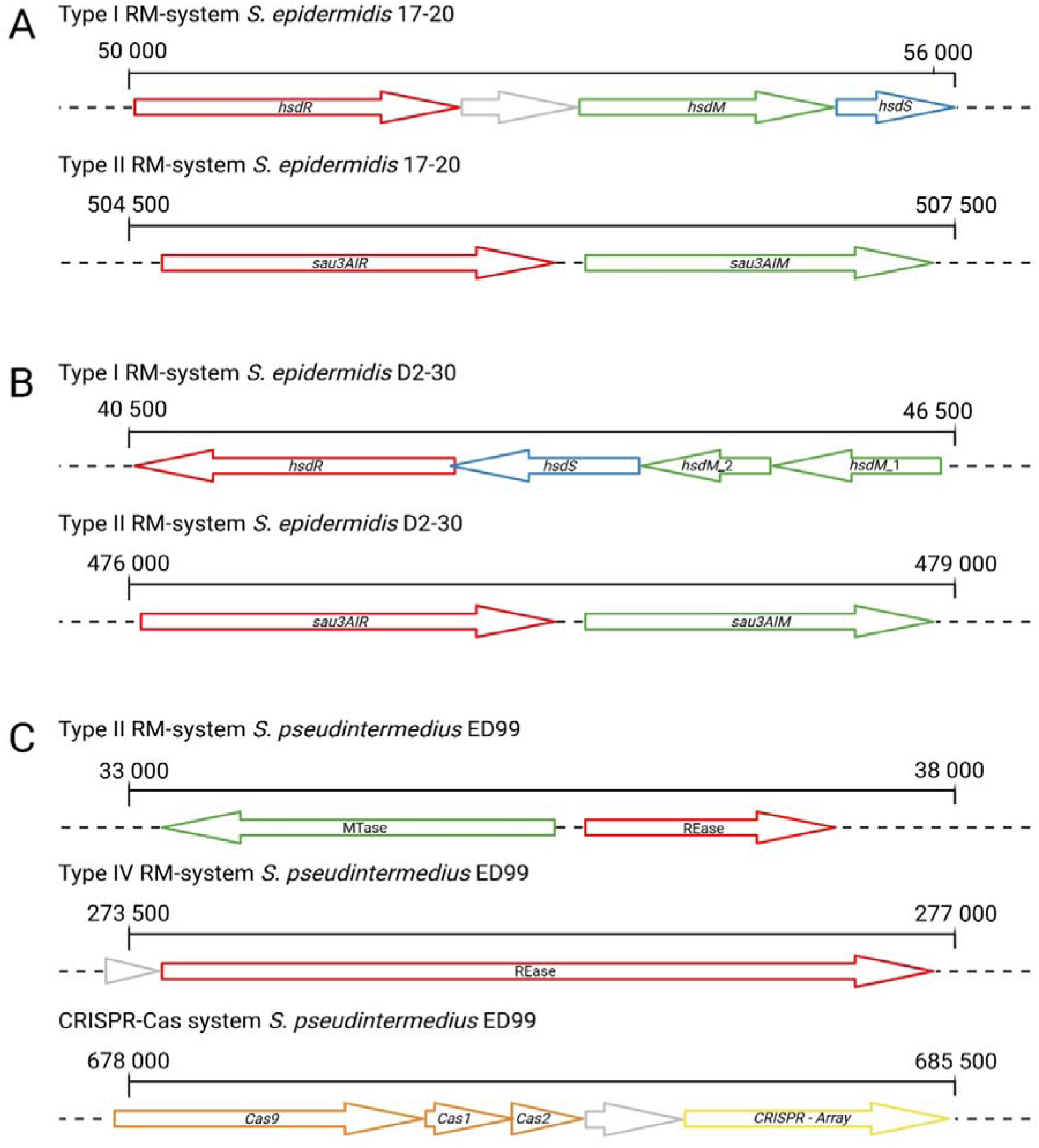
Overview of the RM and CRISPR-Cas systems identified via PADLOC (v2.0.0) for strains S. epidermidis D2-30, S. epidermidis 17-20, and S. pseudintermedius ED99. Methyltransferase genes are highlighted in green; genes mediating sequence specificity of the RM system are highlighted in blue, and genes encoding endonucleases are shown in red. Genes with unknown function are shown in grey. (**A**) Two RM systems were identified for S. epidermidis 17-20. One Type-I system, comprised of the genes hsdR, hsdS, and hsdM. In additon, the cluster contains a gene (EJFLOGOG_00047) with unknown function. The other system is the Type-II sau3AI system with two genes, for the restriction endonuclease (sau3AIR) and the methyltransferase (sau3AIM). (**B**) Two RM systems were identified in S. epidermidis D2-30. The first is a Type-I RM system, containing the genes hsdR, hsdS, and hsdM. The system contains two hsdM copies (hsdM_1 and hsdM_2). The second system is a Type-II RM system, identical to the sau3AI system shown for S. epidermidis 17-20. (**C**) S. pseudintermedius ED99 possesses a Type-II, as well as a Type-IV RM system in addition to its CRISPR-Cas system. The Type-II system consists of a methyltransferase and a restriction endonuclease gene. The Type-IV system is composed of a single restriction endonuclease gene. The CRISPR-Cas cluster is comprised of three CRISPR-associated (Cas) endonuclease genes; Cas9, Cas1, and Cas2. The cluster also contains a CRISPR array with multiple spacers and a repeat sequence.

*S. epidermidis* D2-30 also encoded one Type-I and one Type-II system (Figure 1 B) with the putative recognition sequences ‘GAAYN_5_TGC’ and ‘GATC’, respectively. The Type-II RM system proteins of the two *S. epidermidis* strains showed 100% identity to each other and to the Sau3AI system proteins of other *S. epidermidis* strains. Similarly, the identity was 70% for the restriction enzyme Sau3AIR and 77% for the methyltransferase Sau3AIM to those of *S. aureus* ^40^, which also uses the GATC recognition sequence.

*S. pseudintermedius* ED99 was found to encode a Type-II RM system with predicted recognition sequence ‘CTRYAG’, a Type-IV restriction endonuclease with unclear specificity, and a CRISPR-Cas array (Figure 1 C). The CRISPR-locus carried a Cas9 endonuclease, a CRISPR-associated nuclease Cas1, a CRISPR-associated nuclease Cas2, and a CRISPR array containing multiple spacers and the repeat sequence ‘GTTTTAGCACTATGTTTATTTAGAAAGAGGTAAAAC’, indicating the presence of a presumably fully functional CRISPR-Cas system. An overview of all relevant defence systems, which may explain the difficulties in transformation of the three strains, is summarized in Supplementary Table 2.

### A brief heat shock renders test strains susceptible to phage transduction

Previously established protocols for electroporation of staphylococci ^41,42^ were not successful for the transformation of the three test strains with different plasmids (see Supplementary Table 3: Overview of plasmids used in this study.). A heat shock, applied to the competent cells prior to electroporation, as previously suggested by Loefblom et al. ^33^, did also not result in any transformants. Since phage transduction is known to be an effective alternative for introducing DNA into staphylococci, we investigated whether the plasmids could be transferred into these strains via phage transduction. Phage ΦE72 was chosen for initial experiments with *S. epidermidi*D*s*2-30 and 17-20, as this phage has been shown to infect bacteria of the species *S. epidermidis* ^43^. Because ΦE72 has also been found to bind to *S. pseudintermedius* ED99^44^, it was analysed for its capacity to transduce ED99. However, no successful transduction occurred in our experiments with any of the three strains, using the standard method.

We reasoned that a heat shock step that has been helpful for improving the yields of electroporation in certain *Staphylococcus* strains might also support the efficacy of phage transduction. To test this possibility, cells of the three test strains were heat-shocked for two minutes at 48°C, 50°C, 52°C, or 54°C prior to the addition of phage lysates. These temperatures were chosen according to the previous report of Loefblom et al. ^33^. We found indeed that the new method generated successfully transduced bacterial cells for all three strains. The significant increase in the transduction efficiency, measured in transductants per plaque forming unit (PFU), was already observed when the cells were heat-shocked at temperatures ≥ 48°C (Figure 2 & Figure 3). However, the heat shock at 54°C led to markedly lower transduction efficiency compared to temperatures between 48°C and 52°C (Figure 2). The optimal heat shock temperature was dependent on the recipient strain and the plasmid used. For *S. epidermidis* 17-20, a temperature of 50°C yielded the highest transduction efficiency for plasmid pRB474 (Figure 2A), while 52°C was optimal for pBTn (Supplementary Figure 1). In the case of *S. epidermidis* D2-30, the 48°C heat-shock led to the highest efficiency with all plasmids (Figure 2B), except pBASE6, for which 50°C was slightly more efficient (Figure 3 & Supplementary Figure 2). Interestingly, little to no difference in transduction efficiency was observed for *S. pseudintermedius* ED99 with all plasmids used between 48°C and 52°C, but 54°C also decreased the transduction efficiency (Figure 2C).

**Figure 2:**
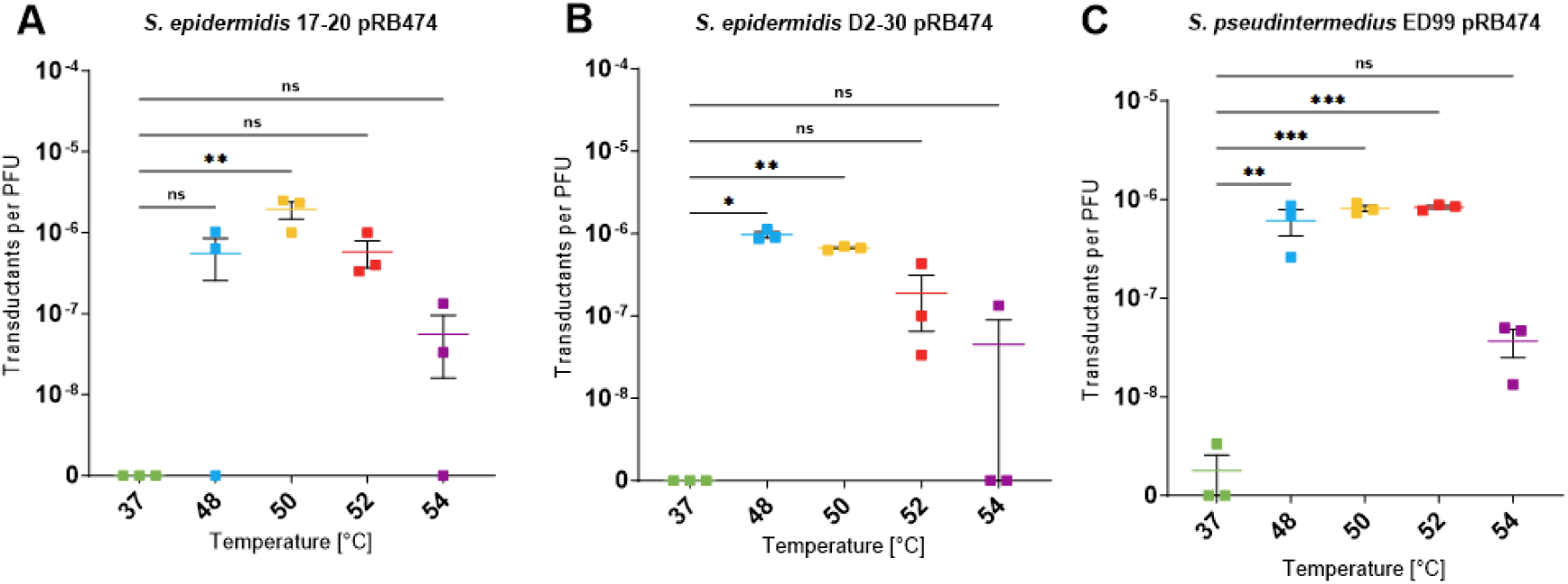
Heat shock transduction of different strains using plasmid pRB474. As control, cells were not heat shocked but incubated for two minutes at the regular growth temperature (**A**) Transduction of S. epidermidis 17-20 wild type. Heat shock prior to transduction led to a marked increase in transductant numbers per PFU, with a peak at 50°C, in comparison to the control (37°C). (**B**) Heat shock transduction of S. epidermidis D2-30. A significant increase in transduction efficiency was observed for 48°C and 50°C. While 52°C still led to a strong increase in efficiency, 54°C nearly abrogated this impact. (**C**) Heat shock transduction of S. pseudintermedius ED99. Temperatures between 48°C and 52°C significantly increased the number of transductants per PFU, whereas 54°C resulted in a loss of transduction efficiency. Data represent the mean of three independent biological replicates (n=3) ± SEM. Statistical analysis was performed via one-way ANOVA. Ns = not significant; *= P < 0.05; ** = P < 0.01; *** = P < 0.001.

**Figure 3:**
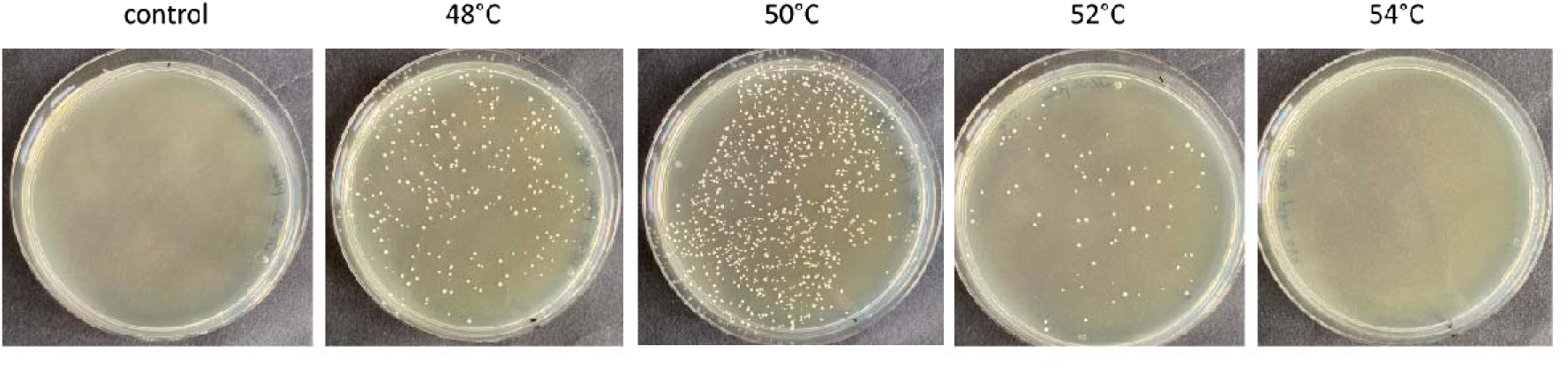
TSA plates, (supplemented with chloramphenicol) 24 h after transduction of S. epidermidis D2-30 with the plasmid pBASE6. No colonies were observed on the control plate, where bacteria were not subjected to heat shock before transduction. However, heat shock at 48°C, 50°C and 52°C enabled successful transduction, resulting in numerous colonies on plates. The highest efficiency was observed at 50°C, whereas heat shock at 54°C generated no colonies. The picture shows a representative transduction replicate.

### The duration of the heat shock influences transduction efficiency

To further optimise the transduction conditions, the heat shock duration was varied between one and ten minutes, using *S. epidermidis* 17-20 at the optimal temperature of 50°C as described above (Figure 2A). We found that our initially chosen duration of two minutes yielded the highest increase in transduction efficiency in comparison to the shorter or longer time periods (Figure 4). A heat shock of one or five minutes still resulted in successful transduction of the strain, although at much lower efficiency, while only residual transductants could be found after the ten-minutes heat shock, indicating that this duration is too harsh for the cells to be efficiently transduced.

**Figure 4:**
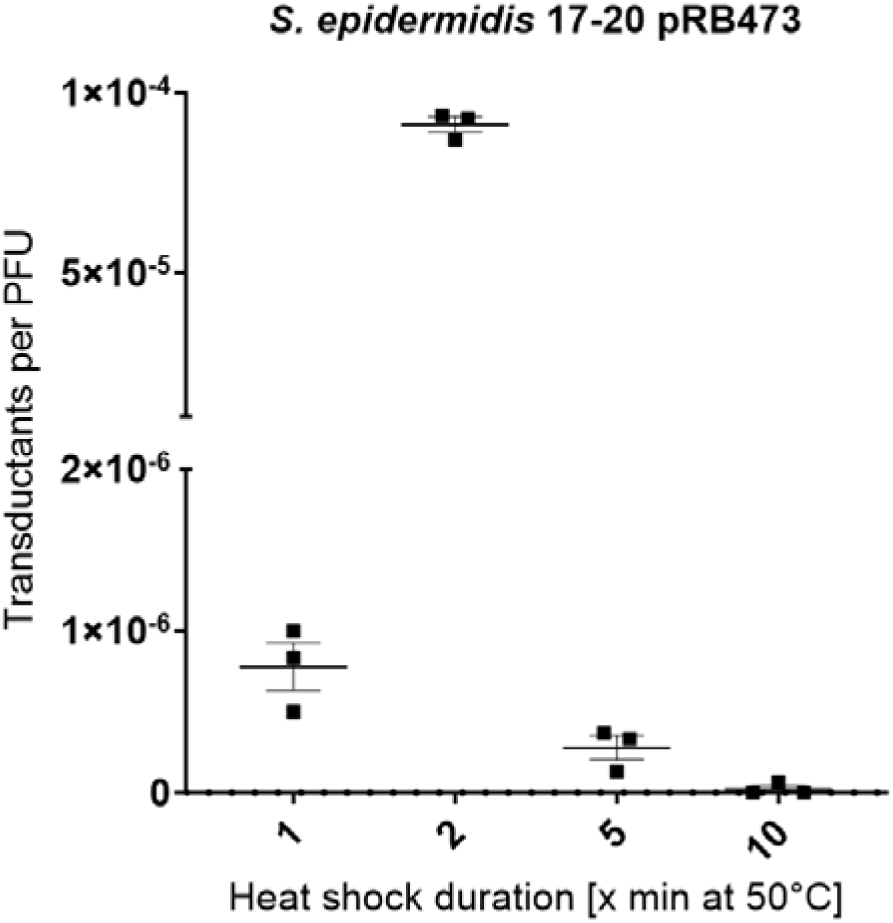
Influence of heat shock duration on transduction efficiency. Wild-type cells of S. epidermidis 17-20 were heat shocked at 50°C for one, two, five, or ten minutes prior to transduction with phage ΦE72 carrying plasmid pRB473. While some clones were growing after one minute and five minutes of heat shock, two minutes was found to be by far the most efficient time span. All data represent n = 3 individual replicates and are shown as mean ± SEM.

### Inactivation of restriction enzymes renders the heat shock prior to phage transduction dispensable

We hypothesised that the restriction systems encoded by the recipient strains are the reason for the inability to transduce the strains with conventional methods, and that the heat shock may have led to temporary inactivation of restriction endonucleases. To test this assumption, we created *S. epidermidis* 17-20 mutants lacking the Type-I RM system (Δ*hsdR),* the Type-II RM system (Δ*sau3AIR),* or both (Δ*hsdR*Δ*sau3AIR)*. The transduction with prior heat shock of the mutant strain panel showed higher transductant numbers, also at the control temperature of 37°C in comparison to the wild type (Figure 5). As observed before, a heat shock of 48°C to 52°C led to a significant increase in transduction efficiency in *S. epidermidis*17-20 wild type, whereas 54°C was less effective (Figure 5 A). While there was still a significant increase in transduction efficiency in *S. epidermidis* 17-20 Δ*hsdR* by the heat shock (Figure 5 B), we found that the heat shock only slightly increased transduction efficacy in *S. epidermidis* 17-20 Δ*sau3AIR* (Figure 5 C) or in the double mutant *S. epidermidis* 17-20 Δ*hsdR*Δ*sau3AIR* (Figure 5 D). The transduction efficiency decreased after the heat shock at higher temperatures probably because it led to damage to the cells.

**Figure 5:**
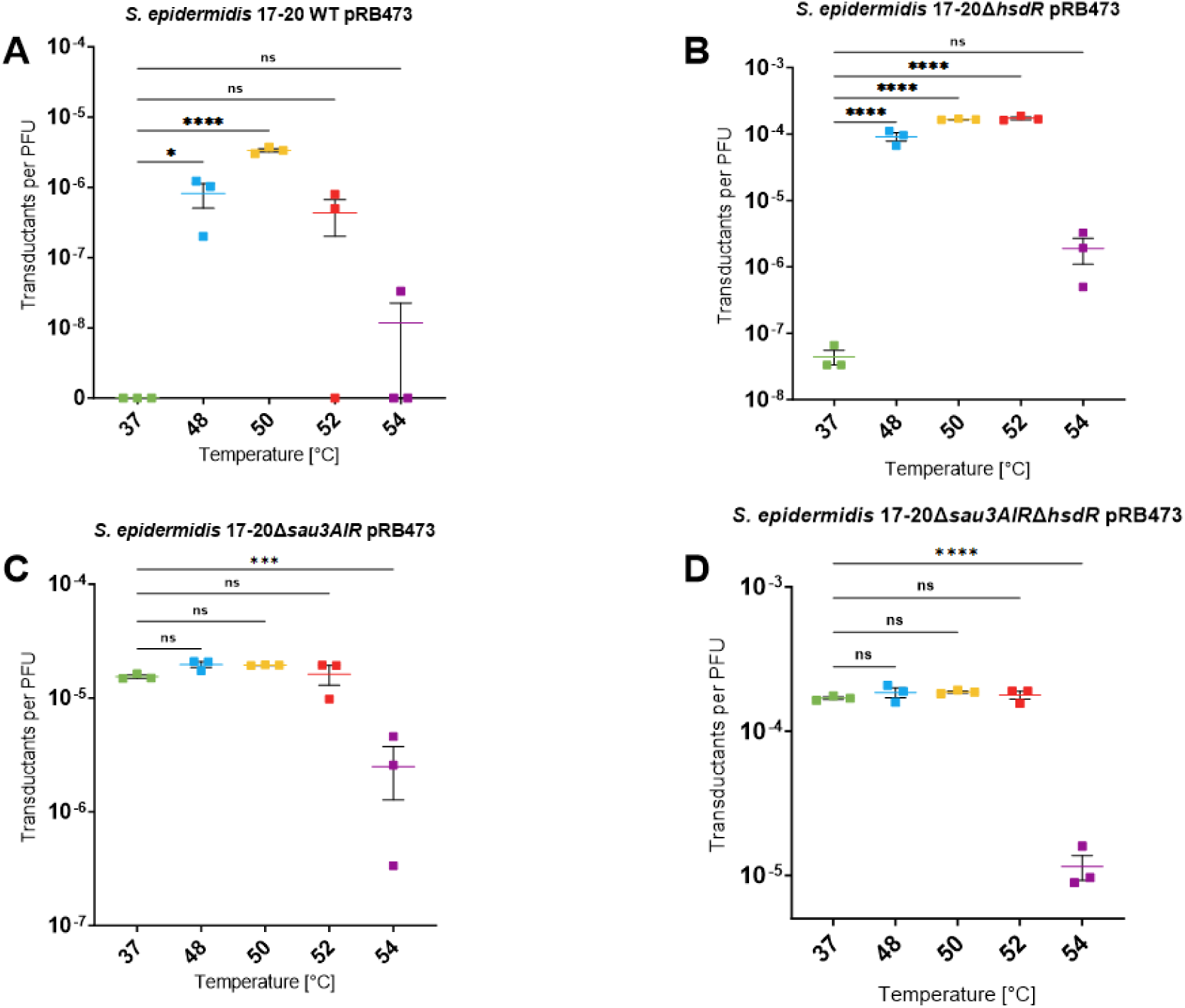
Heat shock transduction of S. epidermidis 17-20 and three restriction-deficient deletion mutants. All experiments were performed using plasmid pRB473. As control, cells were not heat shocked but incubated for two minutes at the regular growth temperature of 37°C. (**A**) Transduction of S. epidermidis 17-20 wild type. Transduction efficiency was significantly increased if a heat shock was applied before transduction. This effect was strongest at 50°C and diminished strongly at 54°C. (**B**) Heat shock transduction of S. epidermidis 17-20ΔhsdR. All temperatures led to a strong increase in transduction efficiency in comparison to the control, however, this effect was markedly decreased at a temperature of 54°C. (C) Heat shock transduction of S. epidermidis 17-20Δsau3AIR. Transduction efficiency at the control condition was higher than in **A** or **B** and there was only a slight increase in efficiency after application of the heat shock between 48°C and 52°C. Heat shock at 54°C led to an overall reduction in efficacy. (**D**) Heat shock of S. epidermidis 17-20ΔhsdRΔsau3AIR prior to transduction did not increase transduction efficacy. However, the heat shock at 54°C led to a significant reduction in efficacy in comparison to the control. All data represent n = 3 individual replicates and are shown as mean ± SEM. Statistical analysis was performed via one-way ANOVA. ns = not significant; *= P < 0.05; ** = P < 0.01; *** = P < 0.001; **** = P < 0.0001.

### Increased transduction efficiency by heat shock is only transient

To analyse for how long the transduction capacity of the test strains persists upon the heat shock, the *S. epidermidis*17-20 wild type was heat-shocked at 50°C and cells were either directly exposed to the phage lysate (t = 0 min) or after 5, 15, 30, or 60 minutes of regeneration in broth at 37°C before transduction. While many transductants were observed at t = 0 minutes, a gradual decrease in efficiency was observed with increasing recovery time. After 60 minutes of recovery, no transductants could be found anymore, indicating that the increased transduction competence was only transient (Figure 6).

**Figure 6:**
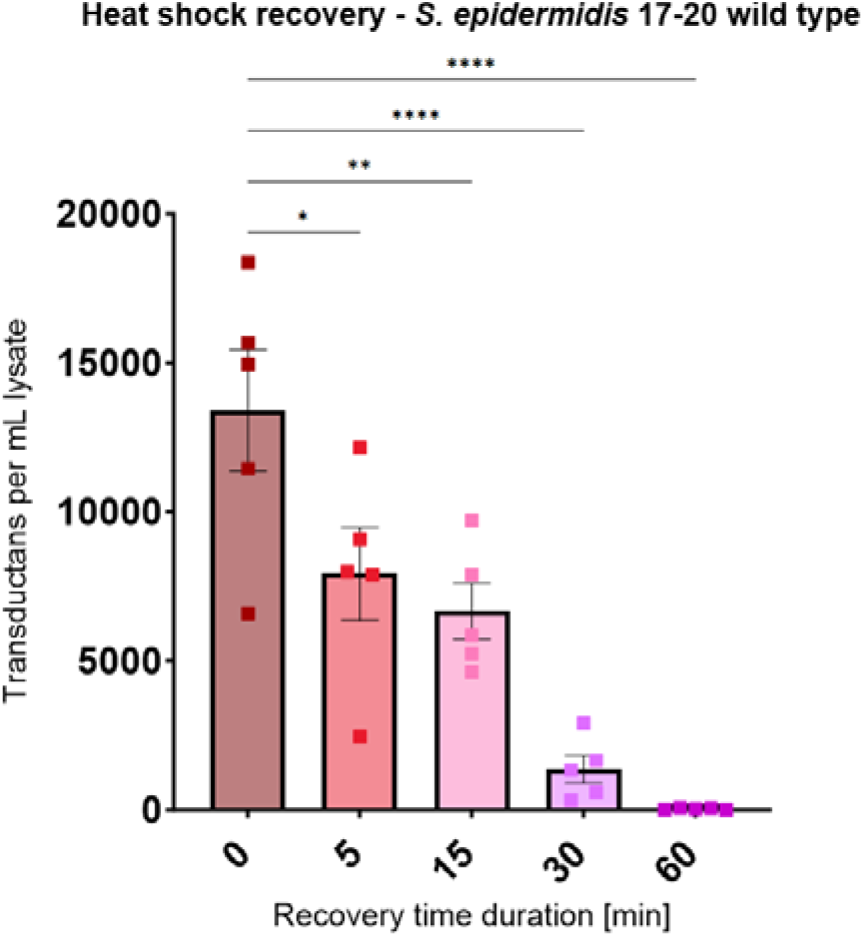
The heat shock recovery assay at 50°C showed a time-dependent decline of the transduction competence. S. epidermidis 17-20 wild-type cells were treated for two minutes at 50°C. After the heat treatment cells were allowed to recover in TSB for 5, 15, 30 or 60 minutes before addition of phage ΦE72 lysate and subsequent transduction (for t = 0 min cells were transduced directly after heat treatment without recovery time to provide a reference value for successful transduction). For the heat shock at 50°C, a time-dependent decrease in transduction efficacy was observed. After 60 minutes of recovery, transduction competence was fully abolished. All data represent n = 5 individual replicates and the mean ± SEM is shown. Statistical analysis was performed via one-way ANOVA. ** = P < 0.01; *** = P < 0.001; **** = P < 0.0001.

### Heat shock increases transduction efficiency also in other bacterial species and genera

We tested the new transduction method on clinical *S. aureus* strains but found no isolates that were difficult to transduce with phage Φ11 propagated on another *S. aureus* strain such as *S. aureus* RN4220. As the transduction rate was already high, the heat-shock protocol did not further improve efficiencies.

To evaluate if the developed protocol for highly efficient transduction is applicable also to difficult-to-transform bacteria other than staphylococci, we studied transduction of plasmid DNA from *S. aureus* RN4220 to *Bacillus spizizenii* and *Listeria grayi,* which also produce ribitol-phosphate wall teichoic acid (WTA), the bacterial receptor structure for the RboP-WTA binding transducing phage Φ11. Both, *L. grayi* and *B. spizizenii*, could be transduced at reasonable efficiency with phage Φ11 carrying the staphylococcal plasmid pT183 even at 37°C without a heat shock (Figure 7). Nevertheless, the application of a heat shock increased the transduction efficiency by up to ∼ 120% or ∼100% (Figure 7 A) in *L. grayi andB. spizizenii,* respectively (Figure 7 B). The two species differed in the optimal heat-shock temperature, which was found to be 54 to 56°C or 50°C for *L. grayi* or *B. spizizenii*, respectively.

**Figure 7:**
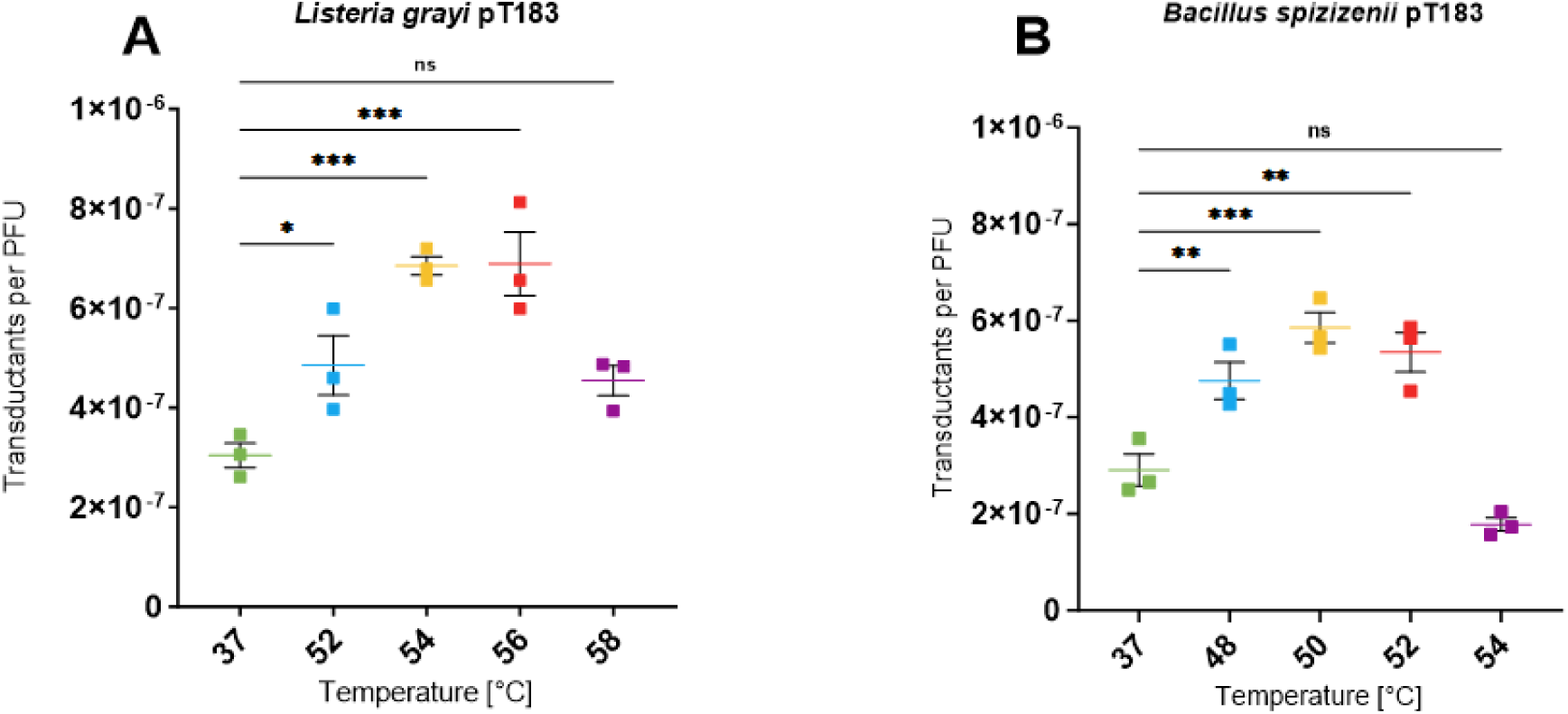
Influence of heat shock on transduction efficiency in Bacillus spizizenii and Listeria grayi using phage Φ11 carrying plasmid pT183. A, Influence on transduction efficiency for Listeria grayi. Cells were heat shocked for two minutes at the indicated elevated temperatures or 37°C as control. While the transduction was possible using the control condition, a significant increase in efficiency was observed at 52°C, 54°C and 56°C, with 54°C and 56°C yielding the strongest increase in efficiency. B, Heat shock transduction of B. spizizenii. Bacteria were heat shocked at the indicated temperatures. Heat shock temperatures between 48 °C and 52 °C led to an increase in transduction efficiency, which was reduced at 54°C. All data are shown as means of three independent biological replicates (n=3) ± SEM. Statistical analysis was performed via one-way ANOVA. ns = not significant; *= P < 0.05; ** = P < 0.01; *** = P < 0.001.

## Discussion

Staphylococci represent key players in the human skin and upper respiratory microbiota, and they have multiple ways to affect health and disease of their host ^4,45–47^. However, while the frequent pathogen *S. aureus* has been studied extensively, the molecular characterization of commensal staphylococci remains challenging. These challenges mainly result from difficulties in genetic manipulation, probably because of the prevalence of one or more foreign-DNA-degrading mechanisms in these bacteria. Classical RM systems have been studied for decades, and the presence of all four types of RM systems has been confirmed in staphylococci ^40,48,49^. More recent studies have found that some staphylococcal clones also encode CRISPR-Cas systems, which may contribute to the barrier, even though they occur in only a minority of staphylococcal isolates ^50–52^.

Since the first discovery of bacterial transduction in *E. coli* by Lederberg et al. in the 1950s ^18^, the adaption of the method for the genus *Staphylococcus* ^19^ and the establishment of DNA transfer to staphylococci via electroporation ^20^, significant advances have been made, enabling the genetic manipulation of various staphylococcal species. These include the optimization of electroporation protocols ^33 42^, identification and use of novel transducing phages ^53,54^, generation of PAM systems ^55,56^, or the use of specifically designed CRISPR-Cas systems^57^. However, many of these methods are only suitable for some bacterial strains or they remain very labour intensive. Despite all these efforts, many *Staphylococcus* isolates remain difficult or even impossible to manipulate. The work presented here aims at providing a method that overcomes transformation barriers even in most refractory strains. Due to its easy, fast, and cost-effective use, we are confident that our method will advance the study of so far inaccessible bacteria, provided that transducing phages are available.

Our experimental data show that a heat shock applied to bacteria prior to exposure to different transducing phages, including the *S. epidermidis* -infecting ΦE72 and the *S. aureus* -infecting Φ11, led to a significant increase in transduction efficiency by, in some cases, multiple orders of magnitude. Application of this heat shock for varying periods significantly influences the efficiency, and two minutes seems to be the most efficient duration for the heat shock in staphylococci. However, it is possible that longer or shorter durations may be more efficient in other bacteria, which needs to be assessed in case-by-case attempts. Notably, this method enabled us to transfer DNA into bacteria that could not be transformed or transduced by any of the previously described standard methods. We also found that a temperature range between 48°C and 54°C is most suitable for heat shock transduction of staphylococci. However, higher or lower temperatures might be more beneficial for the transduction of other species, as found for the transduction of *L. grayi*.

Elevated environmental temperatures are detrimental to cells due to their negative impact on protein stability ^58,59^. We assumed that the heat shock denatures restriction enzymes and therefore enables bacteria to take up and replicate foreign DNA, before degradation by newly synthesized or successfully refolded RM enzymes can occur. We found that both RM systems of *S. epidermidis* 17-20 impacted the ability of strain to acquire foreign DNA, but that the Type-II system, Sau3AI, plays a dominant role, whereas the TypeI system has a subordinate function. However, a reduced transduction efficacy at the highest heat shock temperature of 54°C was obvious. We assume that incubation at this extreme temperature reduces the viability of the cells since temperatures that impair the function of restriction enzymes would also damage other, essential cellular proteins. The cells need to express the plasmid-encoded antibiotic resistance determinant to survive in the presence of the antibiotic but a shift to 54°C could damage various enzymes involved in transcription and synthesis of the resistance-conferring protein. This could in turn force the cells to focus on the protection and re-synthesis of essential cellular components and thereby leaving not enough time to reliably express the antibiotic resistance and as a result also reduce the overall efficiency of the method. This assumption is in accordance with previous findings, indicating that temperatures > 50°C for a short time reduce the viability of staphylococcal cells ^60,61^.

To cope with challenging environmental conditions, bacteria have evolved a large set of fine-tuned response-cascades, including the stringent, cold shock, and heat shock response (HSR), adjusting cellular processes to altered demands ^62,63^. The HSR induces various mechanisms at elevated temperatures involved, for instance, in repair of damaged DNA, stabilization of mRNA, degradation of misfolded proteins, as well as protection of proteins from damage or degradation by chaperones ^63–66^. We observed a gradual decrease in transduction efficiency upon heat shock over time, suggesting that the restriction mechanisms are restored within one hour by the HSR or by de-novo synthesis of restriction endonucleases.

Our heat shock transduction protocol was also effective in other bacterial genera. While *B. spizizenii* and *L. grayi* were also transducible using our standard protocol, a significant increase in transduction efficiency was found after application of a heat shock for both species, albeit at other optimal temperatures as for staphylococci. Thus, our method can be useful for a variety of bacterial species, but the height and duration of the heat shock will need to be optimized for each species.

In summary, our work demonstrates that a short heat shock applied to staphylococci and other bacteria, prior to transduction, significantly increases transduction efficiency and thereby enables the work with otherwise genetically inaccessible strains.

### Limitations of the study

We also found a strong increase in transduction efficiency of *S. pseudintermedius* ED99, which, in addition to a Type-II and a Type-IV RM-system, also encodes a CRISPR-Cas system. It remains to be analyzed if inactivation of this CRISPR-Cas system may contribute to the heat-shock mediated increase of *S. pseudointermedius* transduction.

## Materials and Methods

### Bacterial strains and growth media

All staphylococcal strains used in this work were grown in tryptic soy broth (TSB; Oxoid) except for *S. epidermidis* 1457, which was grown in basic medium (BM, 1% soy peptone, 0.5% yeast extract, 0.5% NaCl, 0.1% K_2_HPO_4_, and 0.1% glucose). *Escherichia coli* was grown in lysogeny broth (LB-Lennox; Sigma-Aldrich) broth. For cultivation on plates, 1.5% agar (15 g L^−1^; Beckton-Dickinson) was added to the respective media, or 0.5% (5 g L-^1^) if soft-agar was prepared. If necessary, plates or liquid media were supplemented with the appropriate antibiotics at concentrations of 10 µg mL^−1^ (chloramphenicol; Sigma-Aldrich), 100 µg mL^−1^ (ampicillin; Carl-Roth) or 12.5 µg mL^−1^ (tetracycline; Carl-Roth). For the preparation of fresh cultures, used for phage transduction or propagation, medium was inoculated with the strain of interest to OD_600_ = 0.1, using overnight cultures incubated for 16-24 h. All liquid cultures were shaken at 140 to 160 rpm and at 37°C. Temperature was decreased to 30°C for incubation of the staphylococci if temperature-sensitive plasmids pBASE6 or pBTn were used. Similarly, plates were incubated at 37°C, except for plates containing staphylococci harbouring temperature-sensitive plasmid pBASE6 or pBTn, which were incubated at 30°C.

### Phage propagation and phage lysate preparation

Phages used in this work were Φ187, ΦE72, and Φ11. Propagation and lysate preparation for phage Φ187 were carried out using the bacterial strain *S. aureus* PS187ΔΔ (Δ*sauUSI*Δ*hsdR)* while phage ΦE72 was propagated in *S. epidermidis* 1457 and phage Φ11 in *S. aureus* RN4220 ^67^.

To propagate the phages, the respective propagation strains were inoculated into liquid cultures and grown overnight (16–24 h). The next day, cultures were diluted to OD_600_ = 0.1 in fresh medium and grown until OD_600_ = 0.4 was reached. CaCl_2_ was added to the culture at a final concentration of 4 mM to increase phage binding. Subsequently, lysate of the phage to be propagated was added (roughly 1/5 of the bacterial culture), and the mixture was shaken at a reduced speed of 70 rpm and 37°C until the culture became clear (2 – 5 h). If no clearing occurred during this timeframe, flasks were incubated overnight at room temperature without shaking, to allow for lysis to occur. Lysed cultures were centrifuged for ten minutes at 4,700 x g to pellet cell debris, and the resulting lysates were sterile filtered using a 0.22 µm filter (Millex®; Merck).

Phage titers were determined by standard plaque formation assay ^68^. To this end, the propagation strain was grown over night in liquid medium. The next day, TSA soft agar (for *S. aureus* PS187ΔΔ and RN4220) or BM soft agar (for *S. epidermidis* 1457) were prepared. After cooling to 50°C, soft agar was inoculated with *S. aureus* PS187ΔΔ, *S. aureus* RN4220 or *S. epidermidis* 1457 overnight culture to OD_600_ = 0.1 and thoroughly mixed, and five mL were poured onto agar plates with either TSA or BM (depending on soft agar used). Plasmid-containing phage lysates or plasmid-free phage propagations were serially diluted to 10^−8^ with phage buffer. 10 µL of the individual phage dilutions were spotted on the soft agar containing the corresponding propagation strain and plates incubated over night at 37°C. The next day, individual plaques were counted at the highest possible dilution, and the phage titer was calculated as PFU mL^−1^.

To prepare lysates for transduction assays, the same procedure was performed, using the propagation strain carrying the plasmid of interest. However, after addition of the phage to bacteria carrying a temperature-sensitive plasmid, the mixture was incubated under shaking at 70 rpm and 30°C instead of the 37°C described above. Propagated phages and lysates were stored at 4°C until use. For transduction, only lysates with a phage titer of at least 5 × 10^8^ PFU mL^−1^ were used to obtain reproducible results.

### Molecular genetic methods

To construct restriction-deficient mutants of *S. epidermidis* 17-20 wild type (Δ*sau3AIR*; Δ*hsdR*; Δ*sau3AIR*Δ*hsdR*), the temperature-sensitive knockout plasmid pBASE6 was used as previously described ^69^. In brief, 1-kb genomic regions directly upstream and downstream of the gene *hsdR* were amplified using the primers KO_hsdR_Up_fwd/KO_hsdR_Up_rev or KO_hsdR_Down_fwd/KO_hsdR_Down_rev, respectively, and digested with the restriction enzymes (Thermo Scientific) indicated in Supplementary Table 1. Fragments were ligated into equally digested pBASE6 using T4-ligase (Thermo Scientific) and introduced into *E. coli* DC10B. The correct assembly of the plasmid named pBASE6_KO_hsdR was confirmed via PCR and sequencing. Subsequently, the plasmid was introduced into the intermediary host *S. epidermidis* 1457 by standard electroporation as previously described ^41^. Lysates of ΦE72 were generated using *S. epidermidis* 1457 pBASE6_KO_hsdR. Plasmids were then introduced via heat-shock facilitated transduction into *S. epidermidis* 17-20 wild type using this phage ΦE72 lysate. The knockout procedure via homologous recombination was performed as previously described, and successful knockout was confirmed via PCR and sequencing using primers KO_hsdR_control_fwd and KO_hsdR_control_rev ^69^.

For the construction of *S. epidermidis* 17-20 Δ*sau3AIR* and Δ*hsdR*Δ*sau3AIR*, the 1-kb upstream and downstream genomic regions of the *sau3AIR* gene, were amplified using primers KO_sau3AIR_Up_fwd/KO_sau3AIR_Up_rev or KO_sau3AIR_Down_fwd/KO_sau3AIR_Down_rev, respectively. The plasmid named pBASE6_KO_sau3AIR was constructed as described above, introduced into *S. epidermidis* 17-20 wild type as well as *S. epidermidis* 17-20 Δ*hsdR* using phage ΦE72, and the homologous recombination procedure was repeated as described above. Successful knockout was confirmed via PCR and sequencing using primers KO_sau3AIR_control_fwd and KO_sau3AIR_control_rev.

### Heat shock transduction assay

All plasmids used for the heat shock transduction experiments with phage ΦE72 were first introduced into *S. epidermidis* 1457 as intermediary host via electroporation or standard transduction using phage Φ187 ^53^. The presence of the plasmids in *S. epidermidis* 1457 was confirmed, and phage ΦE72 lysates were generated as described above. For the transduction assay, recipient strains of interest were inoculated into TSB and gown overnight. The next day, 10 mL of fresh TSB were inoculated with overnight culture to OD_600_ = 0.1. The cultures were grown shaking at 160 rpm and 37°C until OD_600_ = 0.8 was reached. Five 1.5-mL sample tubes were filled with 200 µL of bacterial culture. Cells were centrifuged for one minute at 11,000 x g, supernatant was aspirated, and the pellet was resuspended in 200 µL phage buffer (4 mM CaCl_2_, 1 mM MgSO_4_, 0.1 M NaCl, 50 mM Tris-HCl, 0.1% gelatine (w/v); adjusted to pH 7.8).

One of the aliquots was incubated each at 37°C (control), 48°C, 50°C, 52°C, or 54°C for two minutes in a 1.5 mL thermal block (Eppendorf) without agitation. After two minutes of incubation, 100 µL of plasmid-containing phage lysate was added immediately. Phage–bacteria mixtures were incubated without shaking at 37°C for 10 to 15 min to allow for adhesion and transduction to occur^70^. Afterwards, the mixture was plated on TSA containing an appropriate antibiotic and the plates were incubated over night at 37°C or at 30°C for temperature-sensitive plasmids. If necessary, the mixture was diluted 1:10 with phage buffer bevor plating to allow for colony counting.

After 24h (48 h in case of cells grown at 30°C due to carriage of temperature sensitive plasmids), colonies were enumerated, and the transduction efficiency was calculated as ‘Colony Forming Units’/’Plaque Forming Units’ (CFU/PFU) (Equation 1).

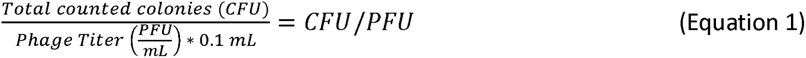

For heat shock transduction of the coagulase-positive species *S. pseudintermedius* ED99, the same procedure was performed. In the case of the species *Listeria grayi* and *Bacillus spizizenii*, phage Φ11 was used. For experiments with phage Φ11, plasmids were introduced into *S. aureus* RN4220 via electroporation ^41^ and lysates with Φ11 generated as described above. Heat-shock transduction was performed as described above for staphylococci, with the exception that the temperature range was increased to 52-58°C for the transduction of *Listeria grayi*.

### Heat shock recovery assay

The assay was performed identically as the heat shock transduction assay described above; the heat shock was applied to the cells for two minutes as well. After incubation of the cells at 50°C or 37°C (control) for two minutes, 800 µL of TSB was added to the cells, and the tubes were incubated at 37°C, under shaking at 160 rpm for regeneration. At t = 0, 5, 15, 30 and 60 minutes of incubation, individual aliquots were centrifuged one min at 11,000 x g. The supernatant was discarded, and the pellet was resuspended in 200 µL phage buffer. 100 µL of ΦE72 phage lysate was added, and the mixture was incubated without shaking at 37°C for 10 min, as described above. Afterwards, the mixture was plated on TSA containing 10 µg/mL chloramphenicol and the plates were incubated over night at 37°C. If necessary, the mixture was diluted 1:10 with phage buffer bevor plating to facilitate colony counting. Transduction efficiency as CFU/PFU was calculated as above using Equation 1.

### Bacterial genome assembly and DNA sequencing

For the assembly of bacterial genomes chromosomal DNA was isolated from the strains and short read sequenced by Illumina while long reads were acquired via Nanopore sequencing. Unicycler v0.5.0 ^71^ with default parameters was used for a hybrid assembly of the Oxford Nanopore and Illumina reads of the *Staphylococcus epidermidis* 17-20 *and* D2-30 genomes. The resulting genome were annotated using prokka v1.14.6 ^72^ with the additional parameters to add gene features in the annotation and searching for non-coding RNAs (parameters --addgenes and --rfam). The quality of the assemblies was assessed using quast v5.3.0 ^73^.

### DNA defence system identification

The Procaryotic Antiviral Defence Locator (PADLOC; v2.0.0 using PADLOC-DB v2.0.0)^38^ was used to identify potential restriction systems encoded in the genomes of *S. epidermidis* 17-20, D2-30 (temporary bioproject number PRJNA1238534, in submission) and *S. pseudintermedius* ED99 (Accession number CP002478.1). Assembled genomes of *S. epidermidis* D2-30 and 17-20 were saved in fasta format. Genome sequence of *S. pseudintermedius* ED99 was downloaded from NCBI database in fasta format. Genomes were uploaded to PADLOC webserver and restriction systems evaluated on the site. All searches were performed with the ‘CRISPRDetect’ function enabled to find any potential CRISPR arrays. Confirmation of the identified RM systems was accomplished using REBASE ^39^ while CRISPR-Cas systems were checked via nucleotide and protein BLAST ^74^.

### Statistical analysis

All statistical analyses were performed using GraphPad Prism version 10.0.0 (153). Data were collected in a column table and analysed via One-Way analysis-of-variance (ANOVA) with standard parameters. Multiple comparisons were set to the control column and corrected using Dunnett’s correction. In case of multiple parameters (Recovery assay Figure 6; temperatures and time points) the data was collected in a group table and analysed via Two-Way ANOVA. In this case, multiple comparisons were corrected using the recommended Tukey’s correction. Output style was in all analyses set to ‘GP’ with the following P-values: P ≥ 0.05 = ns = not significant; *= P < 0.05; ** = P < 0.01; *** = P < 0.001; **** = P < 0.0001.

For all transduction assays with data shown as ‘transductants per PFU’, 10^−9^ was defined as limit of detection (LOD) and was defined as zero, as suggested by GraphPad.

## Supplementary Information

**Supplementary Table 1:**
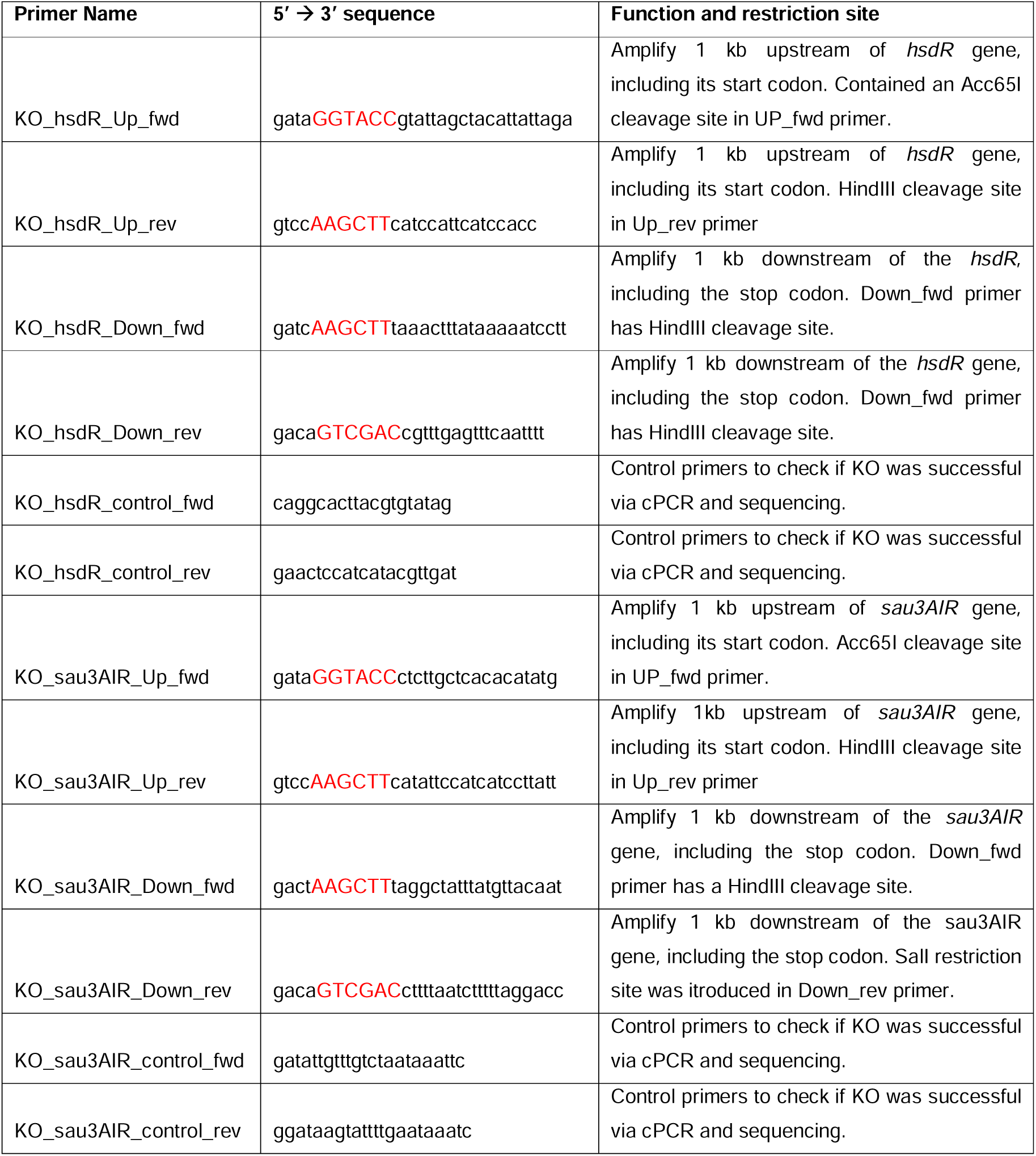
Primers used in this work. If present, cleavage sites for restriction enzymes are written in capital letters and highlighted in red. The corresponding enzyme indicated in ‘Function and restriction site’.

**Supplementary Table 2:**
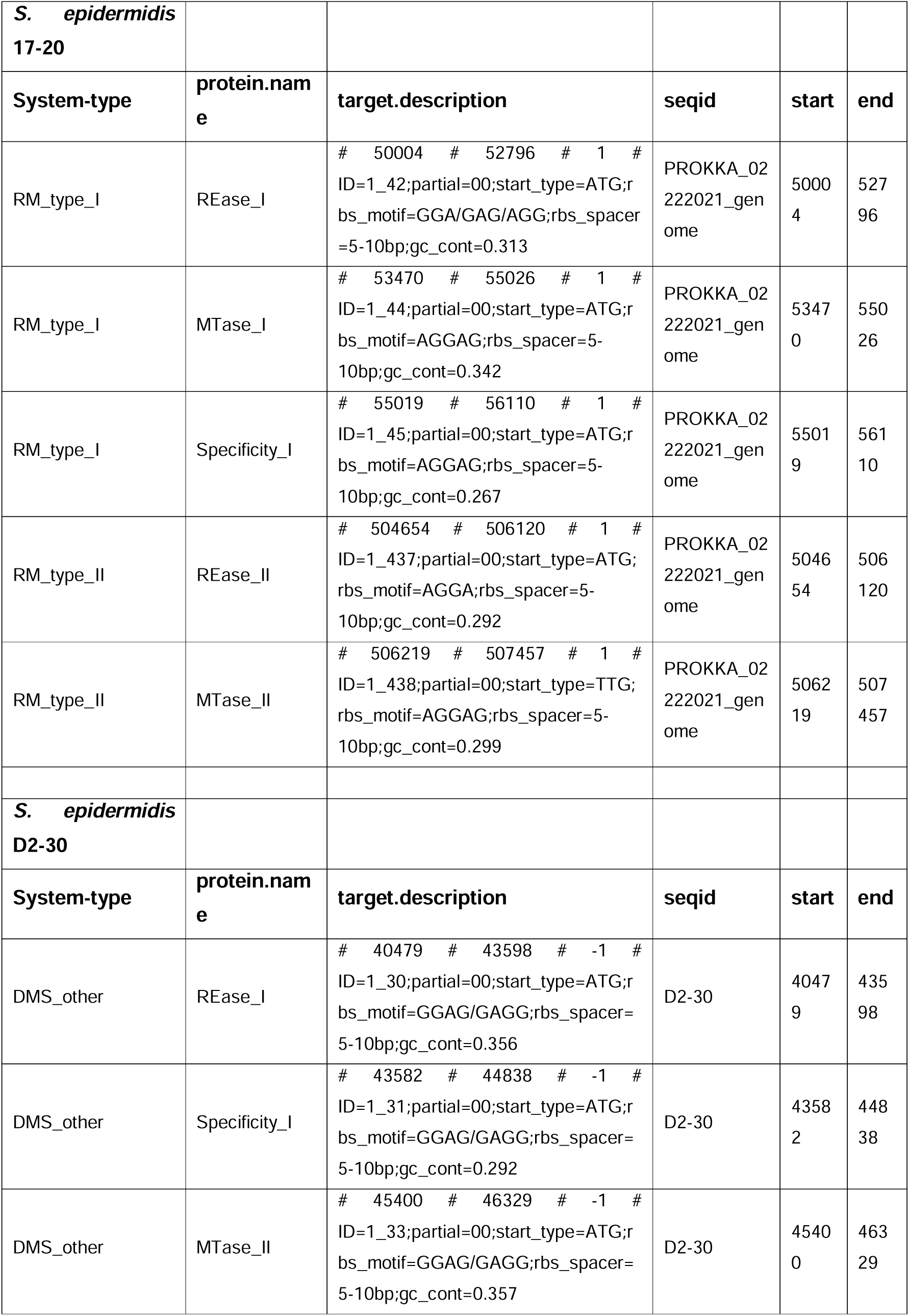
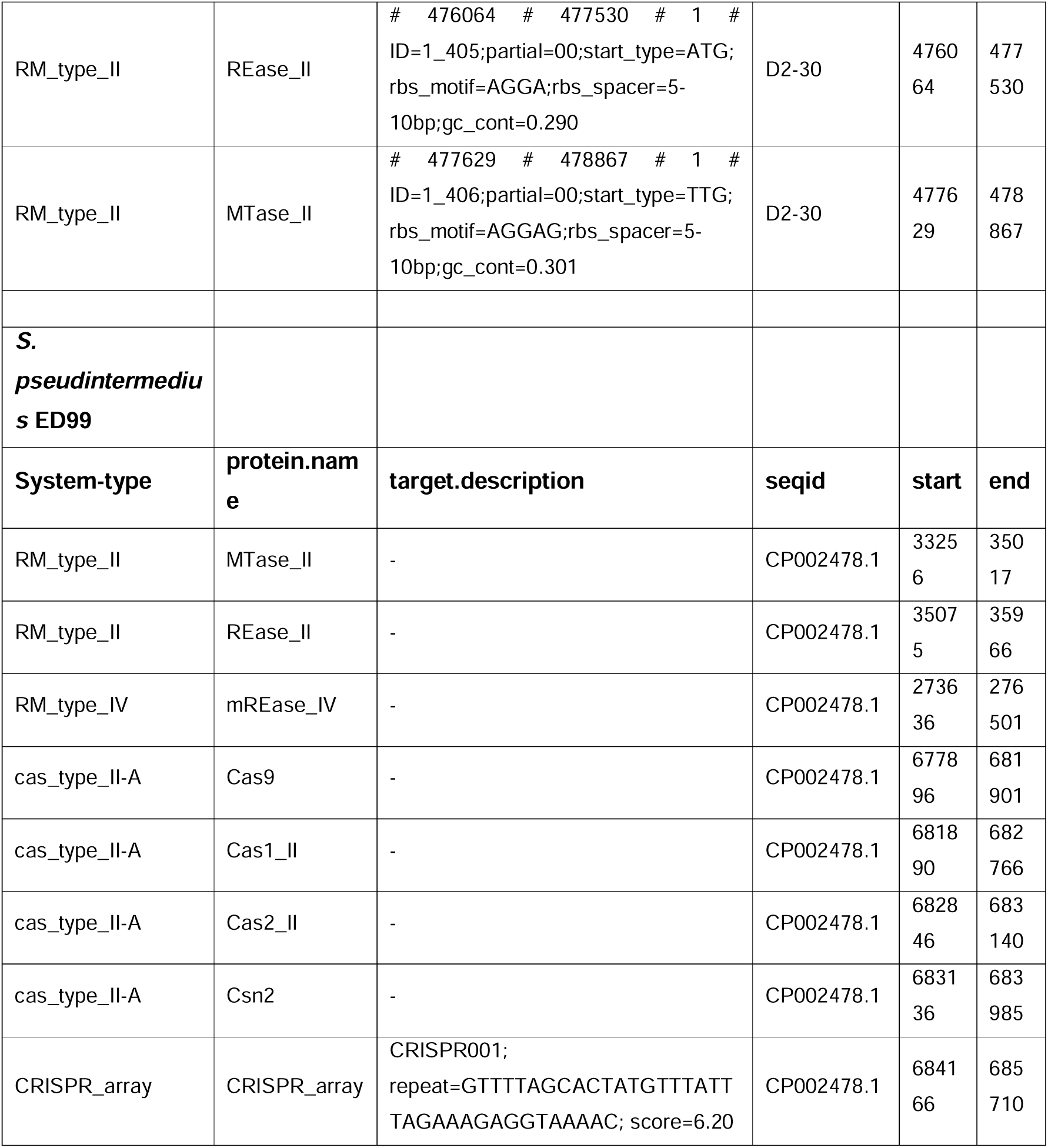
Overview over the different RM systems identified in the strains S. epidermidis D2-30 and 17-20 as well as S. pseudintermedius ED99. Shown are the system type, protein name, target description, sequence ID, and genomic location. All information was derived from the PADLOC webserver.

**Supplementary Table 3:**
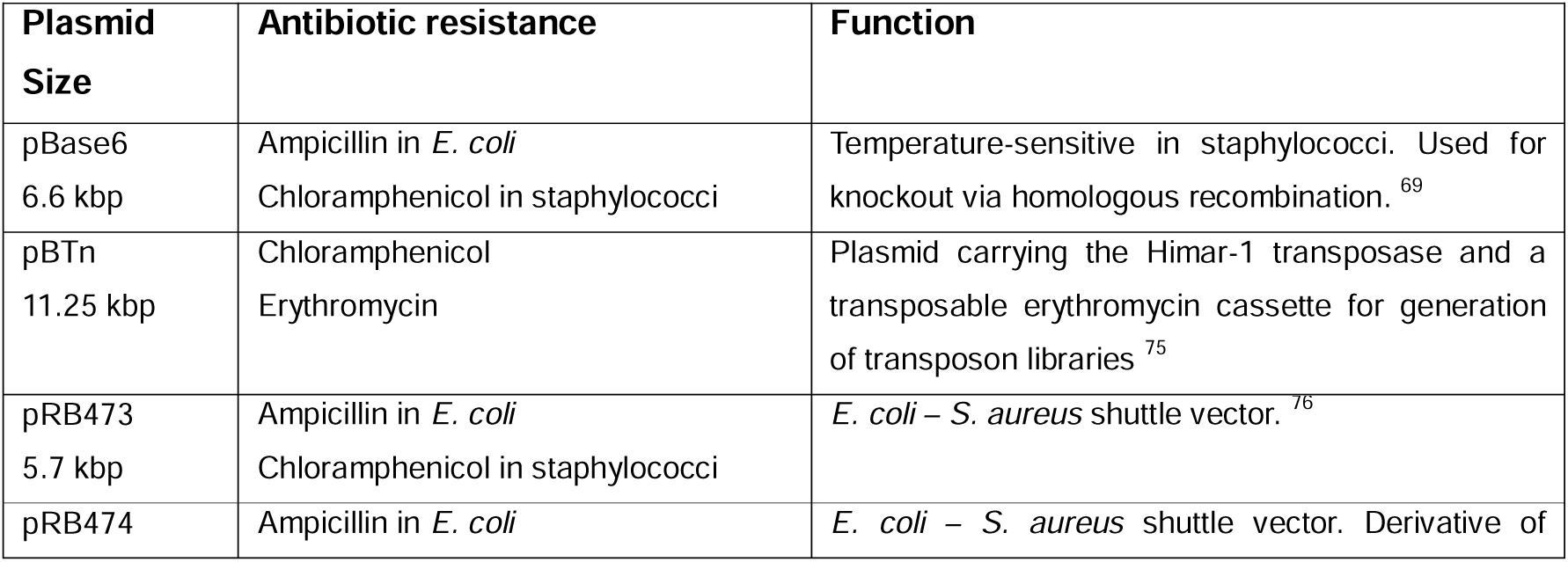
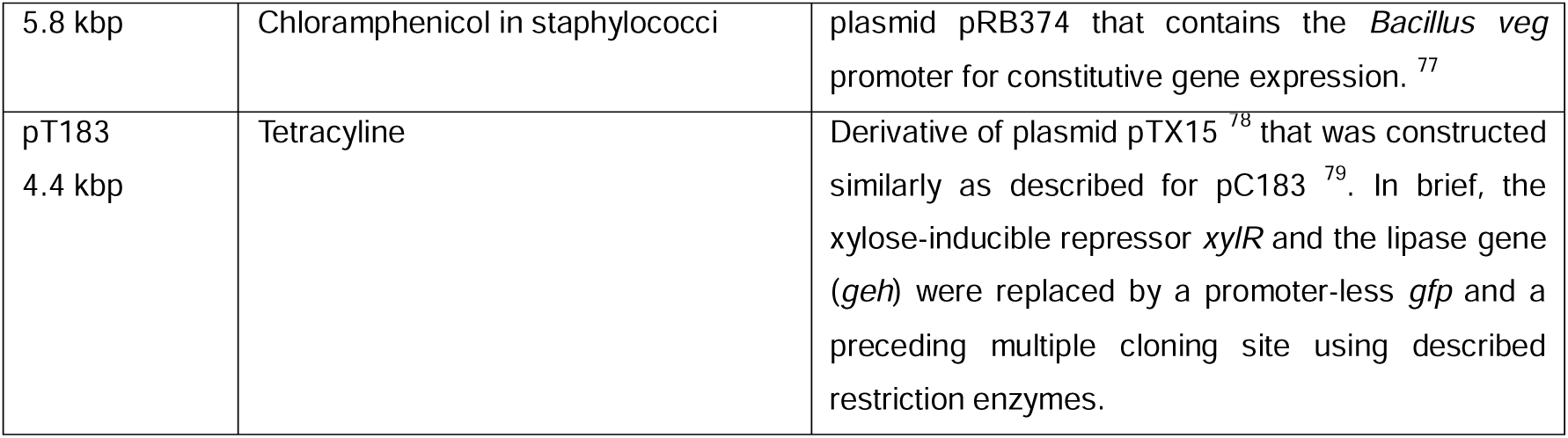
Overview of plasmids used in this study.

**Supplementary Table 4:** Bacterial species used throughout this work.

| Bacterial Species | Description and origin |
| --- | --- |
| *Bacillus spizizenii* W23 | DSMZ strain (DSM 8439). |
| *Escherichia coli* DC10B | $\Delta dcm$ mutant of cloning strain *E. coli* DH10B.<sup>80</sup> |
| *Listeria grayi* | Strain collection AG Peschel, patient isolate. |
| *Staphylococcus aureus* PS187$\Delta\Delta$ | Restriction-deficient variant ($\Delta hsdR\Delta sauUSI$) of *S. aureus* PS187.<sup>67</sup> |
| *Staphylococcus aureus* RN4220 | Restriction-deficient *S. aureus* strain generated by chemical and UV mutagenesis.<sup>67</sup> |
| *Staphylococcus epidermidis* 1457 | Laboratory strain of *S. epidermidis* often used as intermediary cloning host.<sup>81</sup> |
| *Staphylococcus epidermidis* D2-30 | Nasal isolate of *S. epidermidis*. This study. |
| *Staphylococcus epidermidis* 17-20 | Nasal isolate of *S. epidermidis*. This study. |
| *Staphylococcus epidermidis* 17-20 $\Delta hsdR$ | Nasal isolate of *S. epidermidis*, deficient in restriction endonuclease *hsdR*. This study. |
| *Staphylococcus epidermidis* 17-20 $\Delta sau3AIR$ | Nasal isolate of *S. epidermidis*, deficient in restriction endonuclease *sau3AIR*. This study. |
| *Staphylococcus epidermidis* 17-20 $\Delta hsdR\Delta sau3AIR$ | Nasal isolate of *S. epidermidis*, deficient in restriction endonuclease *hsdR* and *sau3AIR*. This study. |
| *Staphylococcus pseudintermedius* ED99 | Clinical isolate of *S. pseudintermedius*.<sup>82</sup> |

**Supplementary Figure 1:**
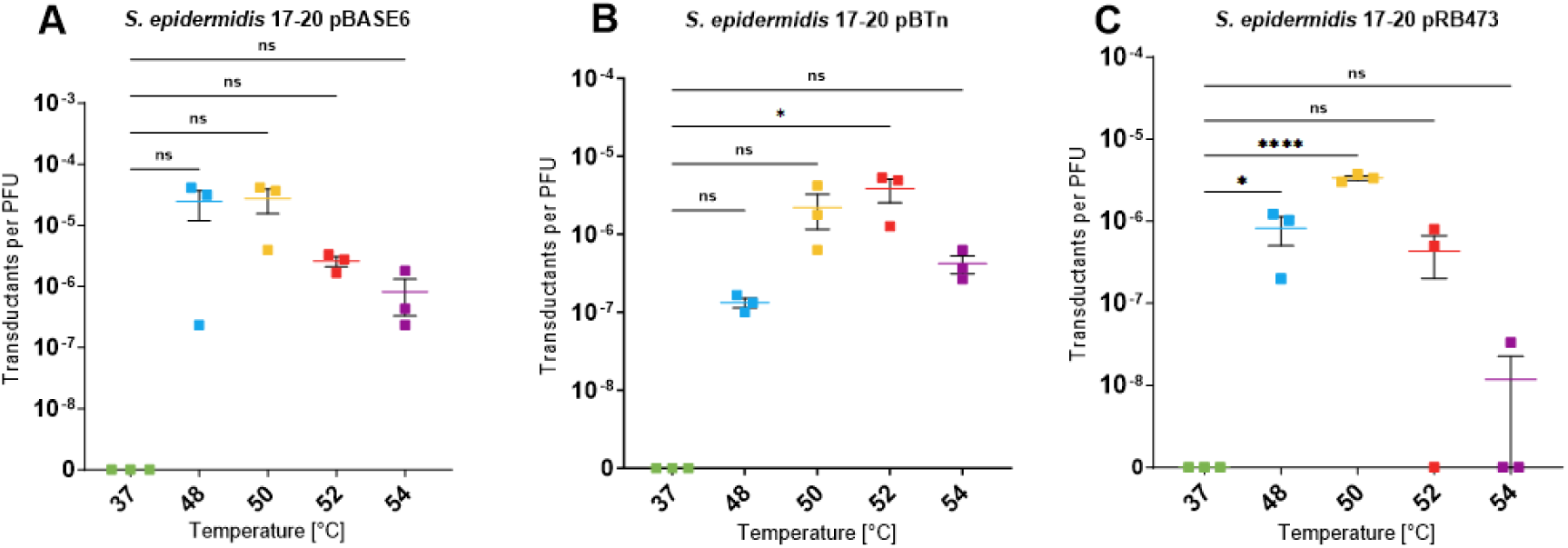
Heat-shock facilitated transduction of S. epidermidis 17-20 using different plasmids. As control, cells were not heat shocked but incubated for 2 min at the regular growth temperature of 37°C. Transduction of S. epidermidis 17-20 wild type with prior heat shock led to a marked increase in transductants per PFU for all three tested plasmids in comparison to the control at 37°C. All data shown as means of three independent biological replicates (n=3) ± SEM. Statistical analysis was performed via One-Way ANOVA. ns = not significant; *= P < 0.05; **** = P < 0.0001.

**Supplementary Figure 2:**
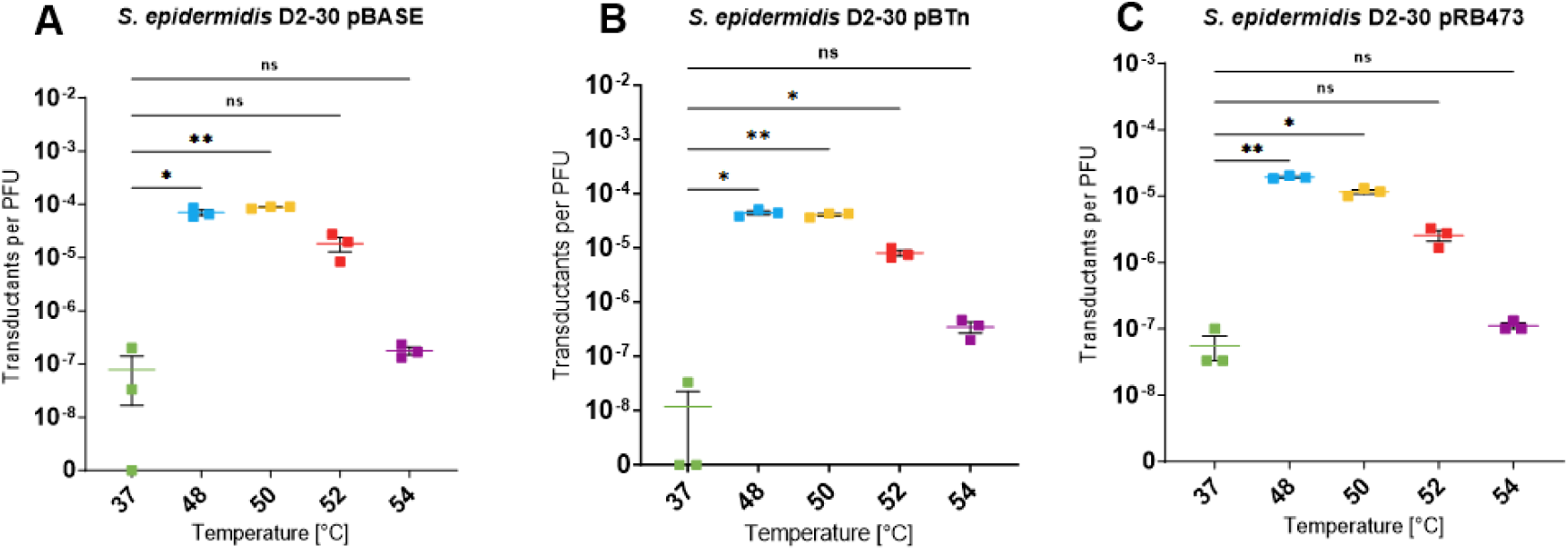
Heat-shock facilitated transduction of S. epidermidis D2-30 using different plasmids. As control, cells were not heat shocked but incubated for 2 min at the regular growth temperature of 37°C. Transduction of S. epidermidis D2-30 wild type with prior heat shock led to a marked increase in transductants per PFU for all three tested plasmids in comparison to the control at 37°C. All data are shown as means of three independent biological replicates (n=3)± SEM. Statistical analysis was performed via One-Way ANOVA. ns = not significant; *= P < 0.05; ** = P < 0.01.

## Resource availability

### Lead contact

Any requests for additional information, mentioned resources or materials should be sent to and will be provided by the lead contact, Bernhard Krismer.

### Material availability

Any material and information used for this study are available from the lead contact upon request.

### Data and code availability

Bacterial genomes were deposited at NCBI and are available under Accession numbers XXX and XXX for *S. epidermidis* 17-20 and D2-30, respectively.

No code has been generated in this study.

Any further data is available from the lead contact upon request.

## Acknowledgements

The authors thank Vera Augsburger for excellent technical support and Janes Krusche and Christian for experimental help. We thank all members of the Peschel and Wolz lab for their help and advice.

## Author contributions

S.K. identified and characterized the strains. K.N. and T.H. sequenced the strains, assembled the genomes and annotated them. B.K. assisted with experimental design, data interpretation and writing the manuscript. J.M. and L.S. designed and performed the experiments, collected and analysed the data and wrote the manuscript draft. A.P. assisted with figure design, data interpretation, manuscript writing and design and finalization.

